# Cost-effective hybrid long-short read assembly delineates alternative GC-rich *Streptomyces* chassis for natural product discovery

**DOI:** 10.1101/2022.12.05.519232

**Authors:** Elena Heng, Lee Ling Tan, Dillon W. P. Tay, Yee Hwee Lim, Lay-Kien Yang, Deborah C.S. Seow, Chung Yan Leong, Veronica Ng, Siew Bee Ng, Yoganathan Kanagasundaram, Fong Tian Wong, Lokanand Koduru

## Abstract

With the advent of rapid automated *in silico* identification of biosynthetic gene clusters (BGCs), genomics presents vast opportunities to accelerate natural product (NP) discovery. However, prolific NP producers, *Streptomyces*, are exceptionally GC-rich (>80%) and highly repetitive within BGCs. These pose challenges in sequencing and high-quality genome assembly which are currently circumvented *via* intensive sequencing. Here, we outline a more cost-effective workflow using multiplex Illumina and Oxford Nanopore sequencing with hybrid long-short read assembly algorithms to generate high quality genomes. Our protocol involves subjecting long read-derived assemblies to up to 4 rounds of polishing with short reads to yield accurate BGC predictions. We successfully sequenced and assembled 8 GC-rich *Streptomyces* genomes whose lengths range from 7.1 to 12.1 Mb at an average N50 of 5.9 Mb. Taxonomic analysis revealed previous misrepresentation among these strains and allowed us to propose a potentially new species, *Streptomyces sydneybrenneri*. Further comprehensive characterization of their biosynthetic, pan-genomic and antibiotic resistance features especially for molecules derived from type I polyketide synthase (PKS) BGCs reflected their potential as NP chassis. Thus, the genome assemblies and insights presented here are envisioned to serve as gateway for the scientific community to expand their avenues in NP discovery.

**Graphic abstract:** Schematic of hybrid long- and short read assembly workflow for genome sequencing of GC-rich *Streptomyces*. Boxes shaded blue and grey correspond to experimental and *in silico* workflows, respectively.

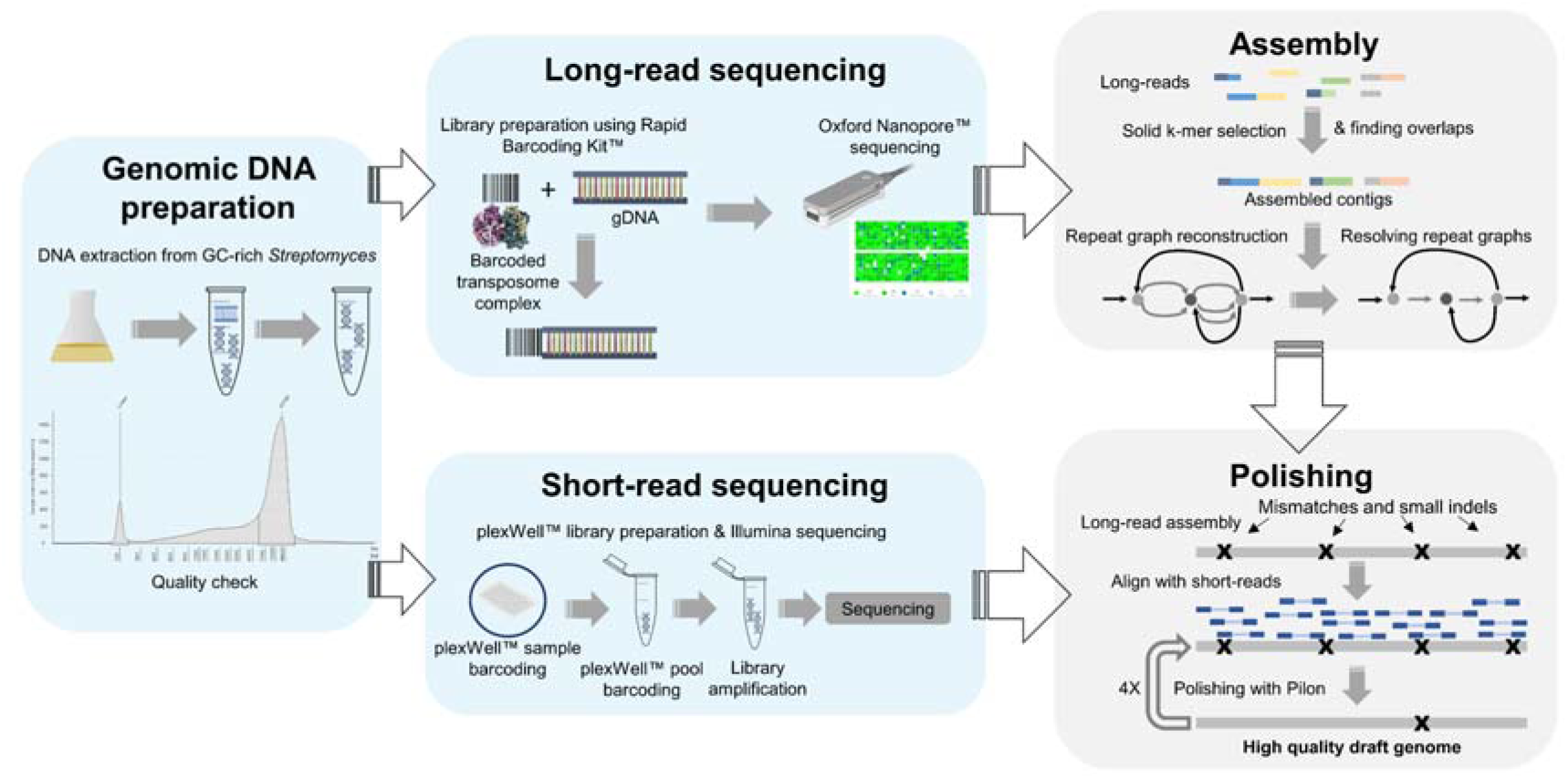

**Highlights:**

- A cost-effective genome sequencing approach for GC-rich *Streptomyces* is presented
- Hybrid assembly improves BGC annotation and identification
- A new species, *Streptomyces sydneybrenneri*, identified by taxonomic analysis
- Genomes of 8 *Streptomyces* species are reported and analysed in this study

## Introduction

Natural products (NPs) are secondary metabolites with myriad applications in therapeutics, agriculture, and food. After decades of technological advancement in deciphering their biosynthetic pathways, we now have highly predictive bioinformatics tools capable of translating genes to biosynthetic information and molecular structures (1,2). This has become vital in the new era of computationally driven antibiotics discovery (3). Due to inactivated gene clusters or low yields of microbial NPs under lab conditions, traditional screening and assay-based drug discovery methods are unable to keep up with the growing crisis of antibiotics resistance and medical needs (3). However, advances in high throughput genomic and bioinformatic tools present a potential reservoir of untapped novel biosynthetic gene clusters (BGCs) in silent strains or in uncultivated microbiomes as sources of new drugs (4).

*Streptomyces* are highly prolific producers and a critical source of NPs, however they are characterized by high-GC content and abundant repetitive sequences, leading to problems in assembling high quality genomes, especially with short-read sequencing (5,6). Instead, long-read sequencing provided by PacBio is greatly preferred to circumvent these limitations (5,6). More recently, Oxford Nanopore sequencing, along with optimized DNA isolation methods, has also been used to obtain long reads for whole-genome sequencing (WGS) of *Streptomyces* (7)(8). While high expense hinders the use of PacBio, Oxford Nanopore sequencing is not exploited to its full potential due to its higher error rate (9). A plausible alternative to this is to employ a hybrid sequencing approach, where assemblies obtained using long reads are subsequently polished with short reads to limit errors in the final assemblies. Here, we describe a simple cost-efficient multiplex sequencing pipeline for the assembly and generation of GC-rich *Streptomyces* genomes. Furthermore, no optimization for multiplex DNA preparation was required beyond commercial protocols for sequencing.

In this study, we were interested in lesser studied *Streptomyces*, including *Streptomyces albulus, Streptomyces diastatochromogenes, Streptomyces lydicus, Streptomyces noursei*, and *Streptomyces* sp. MOE7. With these strains, we demonstrate that standard multiplex Illumina and Oxford Nanopore sequencing protocols can be used in tandem with hybrid long-short read assembly to generate high quality assemblies of lengths ranging from 7.1 to 12.1 Mb and with % completeness varying from 84% for one genome to >93% for the remaining 7 genomes. We further subjected the genome assemblies to taxonomic, pan-genomic and biosynthetic pathway analyses, to resolve misrepresentation in species assignment and explore their potential as alternative NP chassis.

## Results

A total of 8 *Streptomyces* strains were obtained from various sources for this study. 6 *Streptomyces* strains initially annotated as *Streptomyces albulus, Streptomyces diastatochromogenes, Streptomyces lydicus* and *Streptomyces noursei*, were obtained from ATCC (Table 1). *Streptomyces* sp. A44034 and A41733 were obtained from the National Organism Library, Singapore (10) and were annotated as *Streptomyces* sp. MOE7 (NRRL ISP-5461) and *Streptomyces albulus* (ATCC 12757) respectively.

**Table 1.** Whole genome-based taxonomy.

| Strain | Closest Placement Taxonomy | Originally Assigned Taxonomy (Source<sup>a</sup>) | Closest Placement ANI | MSA Percent |
| --- | --- | --- | --- | --- |
| **A41733**<br>(ATCC 12757) | *Streptomyces noursei* | *Streptomyces albulus*<br>(16S) | 97.68 | 92.73 |
| **A44034**<br>(NRRL ISP-5461) | *Streptomyces lydicus* | *Streptomyces* sp.<br>MOE7 (16S) | 99.91 | 89.16 |
| **ATCC11455a** | *Streptomyces noursei* | *Streptomyces noursei*<br>(ATCC) | 99.88 | 87.87 |
| **ATCC21481** | *Streptomyces vietnamensis*<sup>b</sup> | *Streptomyces diastatochromogenes*<br>(ATCC) | 92.93 | 78.49 |
| **ATCC23862** | *Streptomyces diastatochromogenes* | *Streptomyces diastatochromogenes*<br>(ATCC) | 99.88 | 85.05 |
| **ATCC31561** | *Streptomyces mirabilis* | *Streptomyces diastatochromogenes*<br>(ATCC) | 96.6 | 87.65 |
| **ATCC31713** | *Streptomyces albulus* | *Streptomyces albulus*<br>(ATCC) | 99.14 | 81.29 |
| **ATCC31975** | *Streptomyces lydicus* | *Streptomyces lydicus*<br>(ATCC) | 99.91 | 88.74 |

<sup>a</sup>Source refers to how the original taxonomy was assigned; ATCC refers to the assignment as on ATCC (accessed Nov 2022) and 16S refers to assignment by 16S alone.
<sup>b</sup>This is a “low confidence” species assignment by GTDB-Tk (v2.1.0) (11) as indicated by average nucleotide identity (ANI) and multiple sequence alignment (MSA) percent score; the strain is likely to be of different/new species.

### Cost-effective hybrid genome sequencing of GC-rich *Streptomyces*

Long-read sequencing has been especially valuable in sequencing *Streptomyces* whose genomes are characterized by high-GC content and numerous repetitive sequences (5). Although Nanopore-sequenced assemblies are high in contiguity, low per read sequence quality is also common (9). In the context of BGC annotation, this resulting low quality could lead to missing gene or protein domain annotations, thereby resulting in fragmented or mis-annotated gene clusters with incomplete biosynthetic information. Consequently, a combination of long and short reads is required to assemble the vastly repetitive GC-rich sequences of these prolific *Streptomyces*. Prior studies sequencing *Streptomyces* whole genomes have mostly employed PacBio (12) or use one MinION™ flow cell per genome (13)(14), to obtain sufficient coverage and consequently high quality genomes. Here, we demonstrate a simple and convenient multiplexed workflow which does not required further optimization of the genomic preparation for the GC-rich *Streptomyces*. By using standard phenol:chloroform library preparation (15) protocols for these GC-rich genomes; protocols can be performed as described in the commercial kits for multiplex Oxford Nanopore and Illumina sequencing. For Oxford Nanopore, we used the rapid barcoding kit (Oxford Nanopore Technologies, SQK-RBK004) to multiplex 12 genomes on one MinION™ flow cell. For Illumina sequencing preparation, we used plexWell™ 96 (PW096) kit to generate 96 plex samples. The plexWell™ kit uses a reagent limited transposition to ensure uniform read count across samples with various GC and DNA concentrations. Overall cost comparison of Illumina-PacBio sequencing versus our multiplex Nanopore-Illumina protocol estimated that our method of genomic sequencing (including library preparation) has reduced the sequencing costs by >50% (Table S1-2).

### Hybrid assemblies improve genome sequence quality

Nanopore reads are first assembled with the long read assembler, Flye (v2.8) (16), and the resulting assemblies (N50 > 1Mb, Table S4) were polished with short reads obtained from Illumina sequencing (see Methods). Subsequently, we subjected the unpolished and polished assemblies to two different assembly completeness tests: BUSCO (17) and CheckM (18). In our analysis, we observed that the number of polishing rounds was important for assembly quality. Both BUSCO and CheckM analyses revealed a clear improvement in assembly quality following each round of polishing (Figure 1). Maximum improvement occurred after the first round of polishing with subsequent quality improvement almost saturating after 4 rounds.

**Figure 1.**
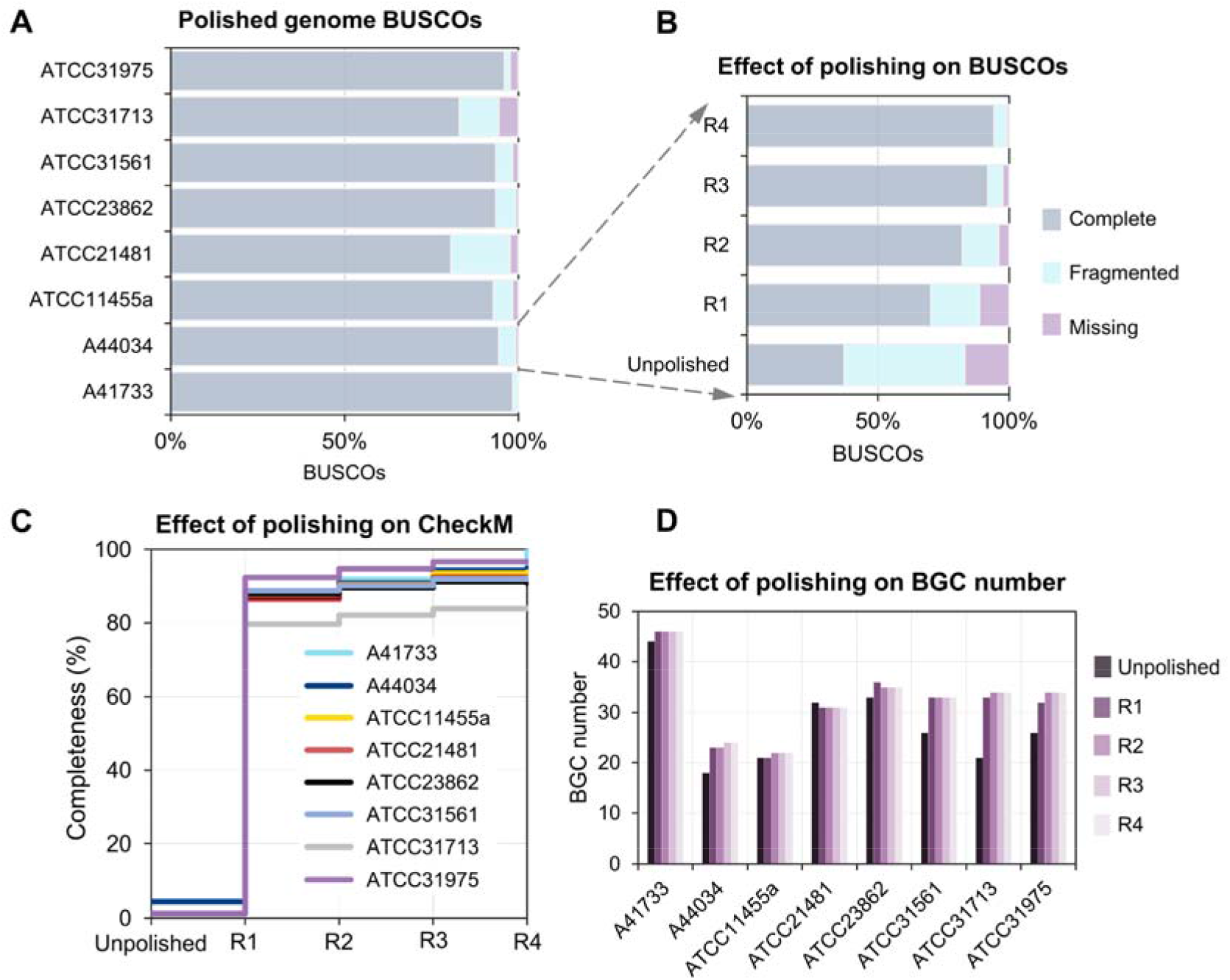
Effect of polishing long read assemblies on genome quality. **(A)** Percentage lineagespecific completeness and contamination levels in polished assemblies as assessed by BUSCO (17). **(B)** Effect of polishing on completeness and contamination. **(C)** Effect of polishing on completeness in polished assemblies as assessed by CheckM (18). **(D)** Effect of polishing on number of BGC identified in assemblies. R1, R2, R3 and R4 in (B-D) correspond to the number of rounds of polishing the long read assemblies with short reads using Pilon (v1.24) (22).

In the 8 high quality genomes assembled for the various *Streptomyces* species, final assemblies observed genome lengths ranging from 7.1 to 12.1 Mb, with an average N50 of 5.9 Mb, and CheckM completeness of 84 to 99% (Table S3). In comparison, the previous genome version of the strain A44034 available on Genbank (accession: GCA_000717345.1) has an N50 of 182 kb. Annotation of the genomes also identified an average of 9850 genes per genome. All except one strain (ATCC 31713) had the completeness values >93 %. The lower CheckM completeness of ATCC 31713 is likely due to its lower quality Nanopore reads (N50 ~70Kb).

Here, we also demonstrate the importance of polishing for sequence quality improvement and its translation to a functional output. Molecular networking of tandem mass spectra (MS/MS) from fermentation extracts of *Streptomyces* sp. A44034 revealed a compound cluster annotated by DEREPLICATOR+ (19) as anti-tumour cyclic siderophore, desferrioxamine E and its rarer analogs (20) (Figure 2A, Table S4). However, to our surprise, antiSMASH (v6.0) of long read assembled (unpolished) and one round of polishing (Figure 2B, Pilon 1) did not uncover a desferrioxamine BGC. Instead, multiple rounds of polishing A44034’s Nanopore assembly with the corresponding short reads were required to achieve sufficient quality to allow accurate BGC annotations. The putative BGC corresponding to desferrioxamine E (21) was observed only after at least two rounds of polishing (Figure 2B, Pilon 2). Also, at Pilon 3, full length genes are now observed for the BGC (Figure 2B). Similarly, we observed an increase in the number and quality of BGCs across the strains with subsequent rounds of polishing (Figure 1D). We therefore highlight the importance of “polishing” to ensure correct and high-resolution analysis of highly GC-rich genomes.

**Figure 2.**
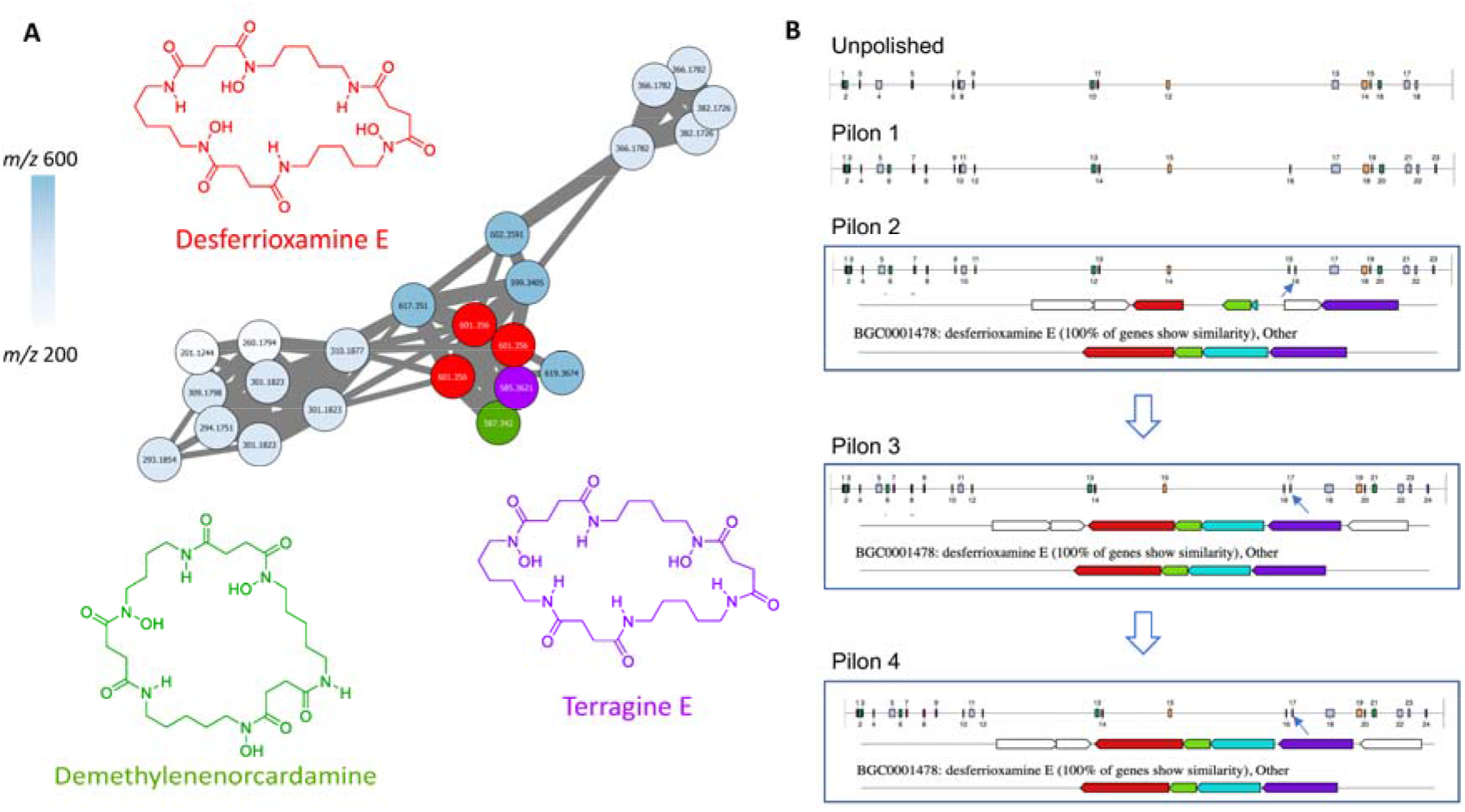
Molecular networking cluster of Desferrioxamine E, and its putative biosynthetic gene cluster (BGC) in *Streptomyces* sp. A44034. **(A)** Desferrioxamine E and its analogs as determined by their MS/MS spectra and molecular networking (23) (Figure S1, S2) **(B)** Revolution of annotated BGCs with increasing polishing rounds. Overview of clusters and their relative positions on the genome, along with multi-gene alignment as analysed using antiSMASH (v6.0) (24) are shown. Blue arrows are used to annotate location of desferrioxamine BGC. Gene cluster table of the putative BGC is available in Table S5. Pilon 1-4 refers to the assembly after 1-4 rounds of polishing, respectively.

### Taxonomy and pan-genomic analysis reveal conserved unique features

All assemblies were subjected to GTDB-Tk based taxonomy (11) (Table 1). Average nucleotide identity (ANI) was used to assign species to the genomes. All except one of the strains (ATCC21481) could be classified into known ‘species’ of *Streptomyces*. ATCC21481 was originally named *Streptomyces diastachromogenes* by American Type Culture Collection (ATCC) at the time of purchase (February 2020). However, our analysis indicates that it has the closest taxonomic proximity to *Streptomyces vietnamensis* (ANI = 92.93). We further subjected ATCC21481 to taxonomic analysis using TYGS webserver (25) and its highest *in silico* DNA-DNA Hybridization (*is*DDH) score was also against *Streptomyces vietnamensis* at 50.8%. Moreover, *is*DDH score between ATCC21481 and *Streptomyces diastachromogenes* (GCA_002242805.1) was estimated to be 22.7%. The current standard species boundary is proposed to be ANI >94 and DNA-DNA Hybridization (DDH) score >70% (26). We further extracted 16S rRNA sequence from ATCC21481 genome and subjected it to BLASTn against the NCBI database. The closest 16S homolog at 99.6% similarity was to an unclassified strain *Streptomyces*, named *Streptomyces sp*. 21-4. Put together, these analyses suggest that ATCC21481 is a potential new *Streptomyces* species within the known Genbank collection. GTDB-Tk based taxonomy (11) (Table 1) was also used to reassign *Streptomyces noursei* to A41733 and *Streptomyces mirabilis* to ATCC31561, which were previously *Streptomyces albulus* (27) and *Streptomyces diastatochromogenes* (28), respectively. The closest 16S homologs to A41733 and ATCC31561 were *Streptomyces noursei* ATCC 11455 at 99.9% similarity and *Streptomyces* sp. RLB1-8 at 100% similarity, respectively.

We then performed pan-genomic analysis of the 8 assemblies along with 4 model *Streptomyces* species using the Anvi’o pipeline v7.1 (29)(see Methods). The pan-genomic map consists of a few distinct regions: the dense region toward the top-left corresponds to the conserved regions most likely harboring genes involved in housekeeping and other core functions (Figure 3). We mapped the presence of the 5 most commonly studied housekeeping genes, rpoB, gyrB, atpD, recA and trpB (30). These were all positioned inside this dense ‘Core’ pan-genomic region. Significant segments contributing to the biosynthetic diversity in different strains also are observed in regions that are unevenly spread across the strain genomes. Interestingly, the accessory regions (‘Cloud’ and ‘Shell’ pan-genome) of the ATCC genomes were enriched with type I polyketide synthases (PKSI) (bottom left of Figure 3). The regions annotated with PKSI were comparatively less in the prototypical *Streptomyces*. A detailed analysis of different BGC classes is presented in the next section.

**Figure 3.**
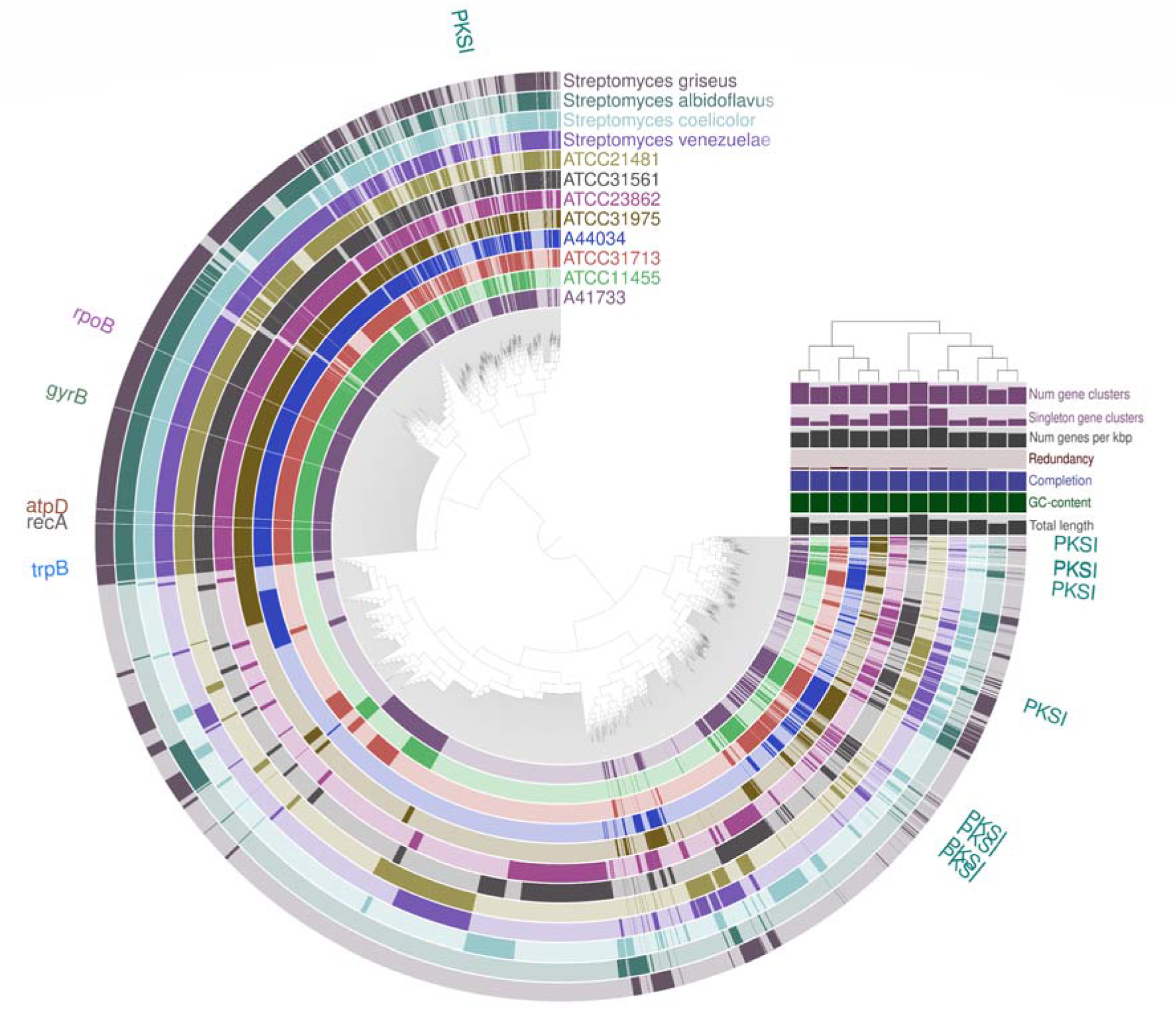
Pan-genomic analysis of the *Streptomyces* strains. The circular strip represents the presence/absence of gene clusters in each of the strains. PKSI: type I PKS; rpoB, gyrB, atpD, recA and trpB are housekeeping genes of *Streptomyces*. Bar plots at the end of each strip represents additional layers summarizing pan-genomic statistics of strains. ‘Num gene clusters’ represents the total number of gene clusters in strains, ‘Singleton gene clusters’ are orphan clusters which do not have homologs in other strains. Dendrogram is constructed using gene cluster presence/absence and Ward clustering. Tree over the layers is constructed based on Euclidean distance between the corresponding values for the strains using hierarchical clustering. Numbers corresponding to different layers are provided in detail in Table S6.

### BGC repertoire and gene cluster family diversity

With genomes of 8 strains assembled, we also sought to explore the BGC diversity among them. To compare if these strains encode any unique BGCs compared to prototypical *Streptomyces* species, we also included *Streptomyces coelicolor* (Accession: AL645882.2) (31), *Streptomyces griseus* (Accession: NC_010572.1) (32), *Streptomyces albidoflavus* (Accession: NC_020990.1) (33), and *Streptomyces venezuelae* (Accession: NZ_CP018074.1) (34) to our analysis on BGC diversity comparisons. First, we constructed a whole genome-based phylogenetic tree using M1CR0B1AL1Z3R (35) and iTOL (36) (see Methods). AntiSMASH (24) based BGC annotation was then performed on all strains. The identified BGCs were then subjected to BiG-SCAPE (v1.1.0) (37) based annotation of classes and gene cluster families (GCFs) while including BGCs from MiBIG 3.0 (38) in the analysis. Finally, the BiG-SCAPE-based BGC classes were aligned to the phylogenetic tree (Figure 4). Clearly, we observe that these strains encode more type I PKS BGCs compared the model *Streptomyces*, possibly suggesting that they could be better hosts than the model *Streptomyces to* produce PKSI-derived natural products. Within A41733, A44034, ATCC 23862, ATCC 31975, we observed desferrioxamine E and a derivative of streptolydigin (Table S4). Although this is only a fraction out of the gene clusters identified *in silico* including pathways towards curamycin, quinomycin, lantipeptide planosporicin, citrulassin D and nystatin (Table S8, S9, S12, S15), it is not unexpected and can be contributed to factors such as unfavourable fermentation conditions (39) or silent gene expression (40). Within these 8 genomes, 27 clusters with less than 30% homology to known MiBiG 3.0 clusters were also observed as potentially new unique BGCs (Table S8-S15). Furthermore, we used antibiotic resistance target seeker (ARTS) (41) to identify BGCs with potential bioactivity. We were able to identify the presence of two categories of resistance, namely the ‘known-resistance mechanisms’ and ‘core duplications’ among multiple BGCs (Figure 4A and 4B). Most strains, except for ATCC 21481 and ATCC 31713, show multiple resistance mechanisms as part of their BGCs. We observed increased presence of ABC efflux transporters among the known resistance mechanisms, indicating potential bactericidal activity among the BGCs (Figure S3). It is interesting to find 1-deoxy-D-xylulose-5-phosphate synthase (*dxs*, (42)) duplicated in the terpene BGCs, as 1-deoxy-D-xylulose-5-phosphate is also intermediate in the non-mevalonate pathway required for terpenoid biosynthesis in *Streptomyces*. However, further experimental validation, and perhaps even strain engineering to activate these clusters, is required to determine if these are indeed critical duplications to combat bioactivity of the corresponding biosynthetic product in the gene clusters.

**Figure 4.**
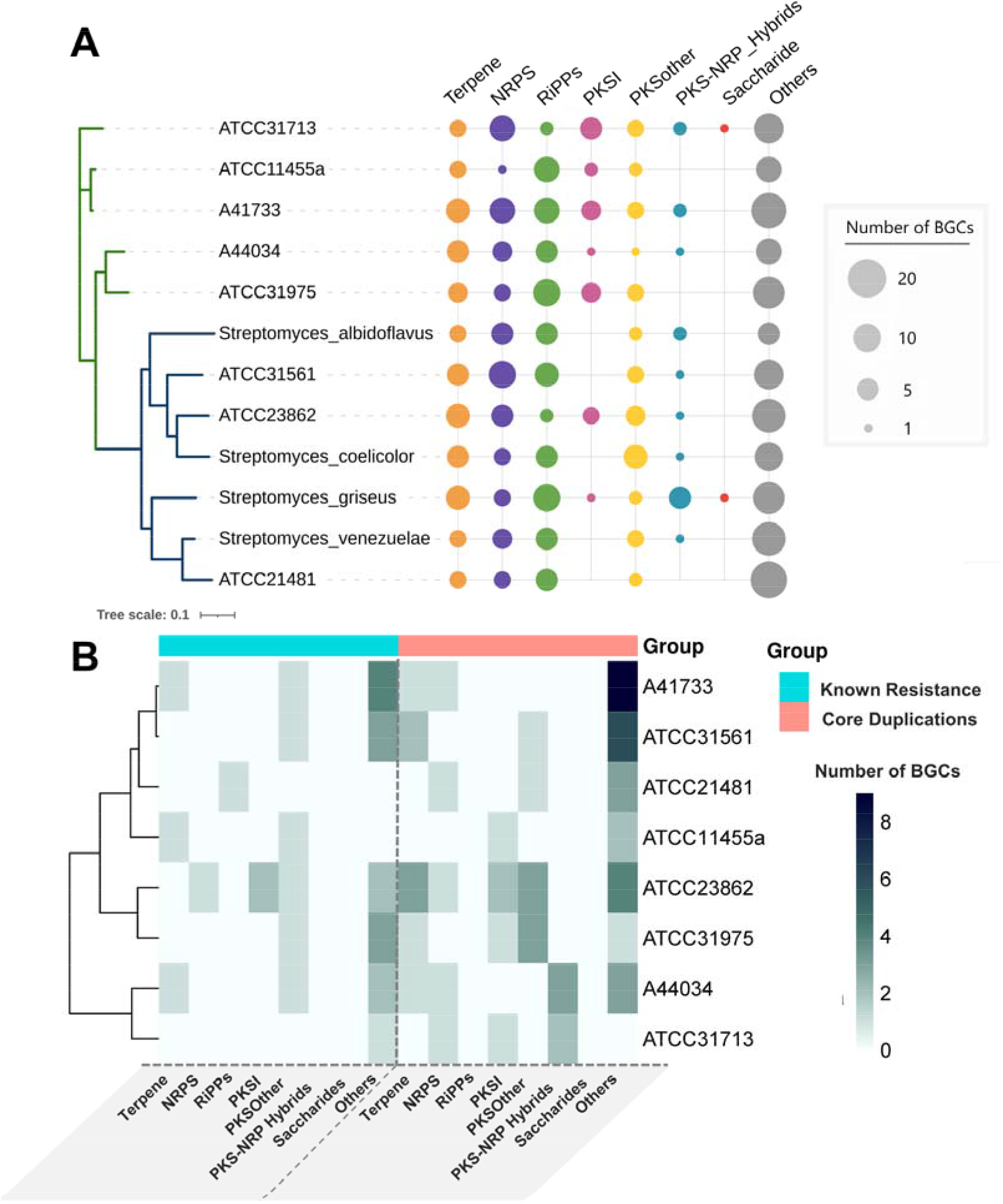
Mining BGCs and resistance mechanisms. **(A)** Enrichment of BGC classes in different *Streptomyces* strains. The size of the circle is proportional to number of BGCs harboured by the corresponding strain. The tree is constructed based on whole-genome phylogeny (bootstrap=100). The clade overrepresenting model *Streptomyces* is coloured blue, while the strains in this study mostly fall under the green clade. More details are provided in Table S7. **(B)** Presence of known-resistance mechanisms and duplicated core genes inside BGCs of different classes as identified by ARTS analysis (v2.0) (41). NRPS: Non ribosomal peptide synthase, RiPPs: Ribosomally synthesized and post-translationally modified peptides, PKSI: Type I PKS, PKSother: PKS other than type I PKS, PKS-NRP_hybrids: Hybrid PKS-NRPS, Others: Cluster that does not fit into any of categories shown.

In addition, we carried out a biochemical network-based pathway enrichment analysis to assess the strain metabolic capacities. The four prototypical *Streptomyces* species and an outgroup heterologous host, *Escherichia coli*, were included in the analysis to facilitate a thorough comparison. The procedure has been detailed in Methods section. Briefly, we first performed protein annotations on the genome assemblies of all strains using Bakta (v1.4.0) (43), the annotations were then used to build draft metabolic networks using Chimera pipeline (44). Finally, KEGG pathway information was extracted from files containing the genome-wide metabolic mapping information (Figure 5). It is interesting to note that two distinct clades emerge from the dendrogram: clade-2 is significantly more enriched in biosynthetic pathways related to several NPs compared to clade-1 (p-value = 1.2055e-04, chi-squared = 14.78, Kruskal-Wallis test). Clade-2 also include commonly used *Streptomyces* chassis, golden standard *Streptomyces coelicolor* (31) and fast-growing *Streptomyces venezuelae* (34). Also, 5 out of the 8 strains examined in this study, including A41733, ATCC23862, ATCC31561, ATCC31713 and A44034, belong to clade-2, hinting at the potential of these strains as alternative chassis for NP production.

**Figure 5.**
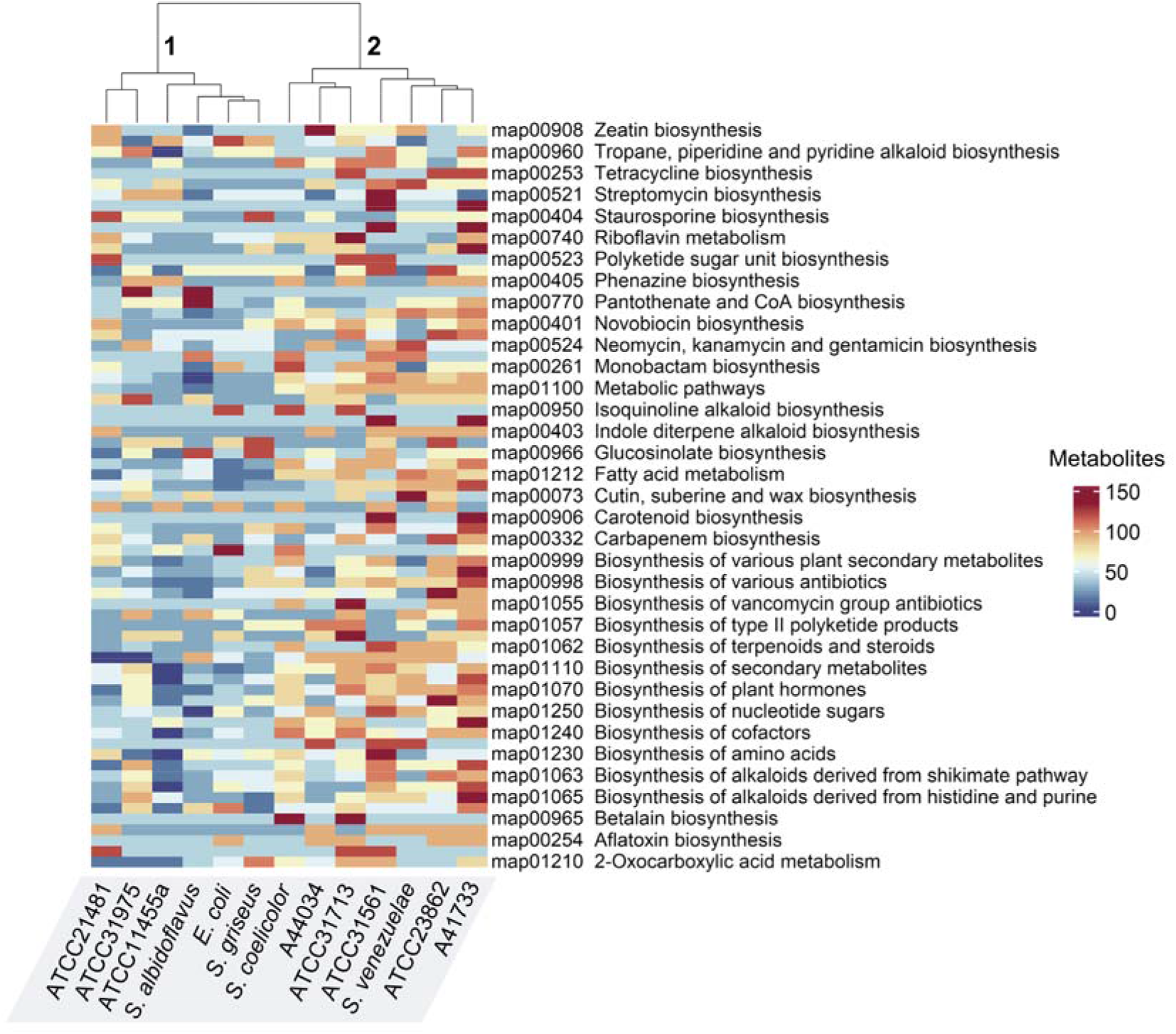
Biochemical network-guided metabolic pathway enrichment. The heatmap showcases enrichment of different KEGG pathway maps related to NP biosynthesis in the 8 strains, 4 prototypical *Streptomyces* and an outgroup heterologous host, *E. coli*. The color scale corresponds to number of metabolites in each KEGG pathway, red being ‘high’ and blue corresponding to ‘low’ values. Labels ‘1’ and ‘2’ at the top represent two distinct clades showing significant differential enrichment in NP biosynthetic pathways (p-value = 1.2055e-04, chi-squared = 14.78, Kruskal-Wallis test). The ‘map’ IDs denote KEGG pathways (45). The dendrogram was generated using hierarchical clustering. Detailed numbers can be found in Table S16.

## Discussion and conclusions

Drastic cost reduction of NGS technologies has greatly accelerated accessibility to genome sequencing, consequently resulting in new applications for exploitation of microbial genomes (46). However, the field of natural product discovery also relies on extensively curated genome sequences with high contiguity (47), which allows for better annotation of BGCs due to accurate positional context of various genes in the genome. Oftentimes, microbes of interest, such as actinobacterial NP producers, are GC-rich and repetitive and PacBio and Oxford Nanopore have been the critical drivers of long read sequencing approaches required to generate high quality assemblies. Although Oxford Nanopore sequencing is several orders cheaper than PacBio, the resulting reads are typically characterized by low per read accuracy, especially in GC-rich actinobacteria which are known to be major reservoirs of natural products (5). In this work, we present a multiplexed short- and long-read sequencing protocol to assemble high quality genomes of GC-rich *Streptomyces*. Here, we sequenced 8 *Streptomyces* strains with potential as natural product chassis and assemble them using a hybrid approach. We observe that polishing Nanopore-assembled genomes with Illumina short read sequences significantly improves BGC annotations. With a substantial reduction (>50%) in sequencing costs compared to the more conventional PacBio-Illumina workflow, this protocol has the potential to considerably advance natural product research. Among the 8 sequenced strains, we have also identified one strain (ATCC 21481) as a presumptive new species, which we have renamed as *Streptomyces sydneybrenneri*. Genomic analyses of these strains also suggest that although most of the BGCs were not expressed under laboratory conditions, these strains have significant potential as NP chassis. Pan-genomic comparation of these strains observed that 6 out of 8 strains possess a higher number of type I PKS BGCs compared with model *Streptomyces*, suggesting their potential as better hosts for type I PKS-derived natural products. Genome-scale metabolic model-KEGG mapping also observed an overall enrichment of metabolic pathways related to the biosynthesis of several NPs in 5 out of 8 strains. Most strains also consist of more than one resistance gene/mechanism, implying the existence of putative bioactivity in them.

In summary, this study describes a cost-efficient multiplex protocol that drastically reduces the cost and complexity of generating high quality GC-rich genome assemblies. Here, we use this protocol to obtain genome assemblies of 8 strains, consisting of *Streptomyces noursei, Streptomyces lydicus, Streptomyces diastatochromogenes, Streptomyces mirabilis*, and *Streptomyces albulus*, including a potentially new *Streptomyces* species. Analysis of these genomes indicate high biosynthetic potential, including possible opportunities for their usage as type I PKS chassis or mining of cryptic gene clusters. We also envision that this protocol will aid in accelerate whole genome sequencing and assembly of these traditionally challenging strains, and in doing so, open further opportunities into NP discovery and applications.

## Materials and Methods

### Strains

*Streptomyces* strains (ATCC^®^ 11455a, 21481, 23862,31561, 31713, 31975) were purchased from ATCC Global Bioresource Center (Manassas, VA, USA). *Streptomyces* sp. A44034 and A41733 were obtained from the A*STAR National Organism library (10).

### Fermentation and Liquid Chromatography-Mass Spectrometry (LC-MS)

Seed cultures were inoculated into 50 mL of fermentation media, CA02LB, CA07LB, CA08LB, CA09LB or CA10LB. The cultures were incubated at 28°C for 9 days at 200 rpm shaking. Cultures were then collected, freeze dried and extracted with methanol. The extracts were analysed as described in a previous study (48).

### Genomic DNA extraction

Strains are seeded in 4mL SV2 (1.5% glucose, 1.5% glycerol, 1.5% soya peptone, 0.1% CaCO_3_, pH 7.0) and incubated for 4 days in a 30°C shaker. Cell pellets are collected and lysed with lysozyme at 5mg/mL for 1 hour. Genomic DNA is extracted with equal volumes of phenol:chloroform:isoamylalcohol and precipitated with 0.6M ammonium acetate and ethanol. DNA precipitate was washed with 80% ethanol and reconstituted with 10mM Tris-HCl pH 8.0.

### Quality characterisation

Quality of genomic DNA was verified with NanoDrop™ 2000/2000c Spectrophotometer and Qubit dsDNA BR Assay Kit. Only samples with ratios of 260:280 and 260:230 more than 1.8 and Nanodrop to Qubit ratio of 1.0 to 1.5 were used for Nanopore sequencing. To achieve a ratio of 1.0 to 1.5 for Nanodrop to Qubit ratio, 0.6x Ampure XP beads were used to clean-up fragments lesser than 500bp. The quality of genomic DNA was further validated using Agilent Genomic DNA ScreenTape on a tapestation.

### Genomic library preparation and sequencing

To sequence genomes, a library was generated using 10 ng genomic DNA per strain with plexWell™ 96 kit, as described in its manual. The library was sequenced using 2 x 151 bp paired end protocol on illumina HiSEQ 4000.

400 ng genomic DNA per strain was used for sequencing on MinION™ system (Oxford Nanopore Technologies). Rapid Barcoding Sequencing (SQK-RBK004) kit was used for library preparation. All was done accordingly to provided manual. The library was added to a R9.5 Flowcell for 72 hours run on the MinION™ Sequencer.

### Data processing and genome assembly

Hiseq reads were check for quality using FastQC (49), adaptors were trimmed using Trimmomatic (v0.32) (50). Short reads were assembled using Spades assembler (v3.15.5) (51). Nanopore raw data (FAST5) was basecalled and quality-assessed using Guppy GPU version (Oxford Nanopore Technologies GitHub: Guppy [https://github.com/nanoporetech].). The base called FASTQ reads were assembled using Flye (v2.8) (16) using expected genome size of 8Mb and coverage of 30. Long reads were polished with short reads using Pilon (v1.24) (22). Four rounds of polishing were done to ensure better per read quality. BUSCO (17) and CheckM (18) analysis was performed to check assembly completeness.

### Phylogenetic analysis and taxonomic inference

Whole genome phylogenetic tree was contructed using M1CR0B1AL1Z3R (35). The newick trees were visualized using iTOL browser (36). BGC class bubble plot annotations were added using iTOL’s ‘create dataset’ options. Taxonomic inferences were made based on whole genome assemblies using GTDB-Tk (v2.1.0) (11). A scratch directory was used to avoid RAM limitations of PPlacer algorithm part of the GTDB-Tk pipeline. The *in silico* DNA-DNA Hybridization score (*is*DDH) was estimated using TYGS webserver (25). 16S-based taxonomy was inferred as follows: 16S sequences were extracted from the genome assemblies FASTA files using Barrnap (v0.9) (52)

### Genome mining, BiG-SCAPE, enrichment analysis and resistance guided BGC prioritization pipeline

AntiSMASH (v6.0) (24) in ‘full’ mode was used for BGC annotation. BGC class and GCF annotations were derived from BiG-SCAPE analysis (v1.1.0) (37) with default options while manually including MiBiG 3.0 BGCs (38) to the input. Identification of resistance mechanisms was carried out using antibiotic resistance target seeker (ARTS) pipeline (v2.0) (41). Metabolic pathway enrichment analysis

The genome sequences were first subjected to annotation using Bakta pipeline (v1.4.0) (43). Strain-specific genome-scale metabolic models were generated from the Bakta protein annotations using CarveMe algorithm (v1.4.1) (53) part of Chimera pipeline (44). KEGG (45) pathway information for each strain was retrieved from Chimera outputs and merged to generate pathway enrichment heatmaps. Significant difference in KEGG pathway enrichment between the two dendrogram clades was evaluated using Kruskal-Wallis test. Pan-genomic analysis was performed using Anvi’o (v7.1) pipeline’s default workflow (29).

### DEREPLICATOR+ Metabolite identification

Metabolite identification from tandem mass spectra (MS/MS) data using the online workflow DEREPLICATOR+ (19) available on the Global Natural Products Social Networking (GNPS) website (http://gnps.ucsd.edu, accessed November 2022). Precursor and fragment ion mass tolerance thresholds were set to ± 0.005 Da and ± 0.01 Da respectively. Maximum charge allowed was set to 2. Fragmentation model used was 2-1-3, which indicates there will be at most two bridges, one 2-cut, and three cut in total. The spectra were searched against the “AllDB” small molecule structure database of about 720K compounds.

### Molecular Networking of Desferrioxamine E Cluster

MSConvert v3.0.22198-0867718 from Proteowizard (54) was used for initial processing of raw liquid chromatography-tandem mass spectrometry (LC-MS/MS) data into open-source mascot generic format (.mgf). All tandem mass spectra (MS/MS) signals with intensity values below 1000 signal intensity were removed as background correction. All peaks in a +/- 17 Da around the precursor ion mass were deleted to remove residual precursor ions, and peaks not in the top 6 most intense peaks in a +/- 50Da window were filtered out. MetGem v1.3.6 (55) was used to plot molecular networks with the following filters (m/z tolerance = 0.02, minimum matched peaks = 4, max nearest neighbors = 10, minimal cosine score value = 0.70, max connected component size = 1000).

## Data availability

All genome sequences generated in this study have been deposited to GenBank (Accession: PRJNA907517).

## Author contributions

L.K.: Conceptualization, methodology, software, data curation, formation analysis, visualization, original draft-writing, review and editing, F.T.W.: Conceptualization, investigation, methodology, original draft-writing, review and editing, E.H.: investigation, methodology, data acquisition (genomic sequencing), review and editing, L.L.T.: methodology, data acquisition, V.N., C.Y.L: Data acquisition, strain resources, fermentation, S.B.N.: Data acquisition, strain resources, fermentation, writing-review and editing, L.K.Y., D.S.: Data acquisition, Y.K: Data acquisition, review and editing, D.T., Y.H.L.; Data analysis, writing-review and editing.

## Acknowledgments

The authors gratefully acknowledged the funding for this work from National Research Foundation, Singapore (NRF-CRP19-2017-05-00) and Agency for Science, Technology and Research (A*STAR), Singapore (#21719). The authors also acknowledge the support of the late Professor Sydney Brenner, founder of Molecular Engineering Laboratory at A*STAR, Singapore.

