## Supplementary Information for "Cost-effective hybrid long-short read assembly delineates alternative GC-rich *Streptomyces* chassis for natural product discovery"

**Funding:** This work was supported by National Research Foundation, Singapore (NRF-CRP19-2017-05-00) and Agency for Science, Technology and Research (A*STAR), Singapore (#21719).

**Supplementary information**

**Table S1.** Cost comparison for PacBio versus Nanopore (Singapore based sequencers, Genome Institute of Singapore, A*STAR).

**Table S2.** Cost for Illumina NGS (Singapore based sequencers, Genome Institute of Singapore, A*STAR).

**Table S3.** Genome assembly and annotation of 8 *Streptomyces* strains

**Table S4.** DEREPLICATOR+ metabolite identification

**Figure S1.** MS/MS spectral comparison of desferrioxamine E and demethylenenorcardamine. Cosine similarity = 0.9748.

**Figure S2.** MS/MS spectral comparison of desferrioxamine E and Terragine E. Cosine similarity = 0.8360.

**Table S5.** Desferrioxamine gene cluster from A44034, compared to miBIG BGC0001478: desferrioxamine E biosynthetic gene cluster from *Streptomyces sp.* ID38640

**Table S6.** Numbers underlying different pan-genomic 'layers' in Figure 3

**Table S7.** BGC classes in the *Streptomyces* strains identified using antiSMASH v6.0 and BiG-SCAPE

**Figure S3.** ARTS annotations for Streptomyces BGCs depicted in Figure 4B and 4C. A. Known resistance genes/mechanisms in BGCs. B. Duplicated core genes in BGCs. C. Abbreviations and names of A and B.

**Table S1.** Cost comparison for PacBio versus Nanopore (Singapore based sequencers, Genome Institute of Singapore, A*STAR).

| | **PacBio** | **Nanopore** |
|---|---|---|
| **Library preparation[a, b]** | 2402 | 650 |
| **Sequencer** | 3085/SMRT lane | 555/ R9.4.1 flow cell |
| **Total cost ($USD)** | 5487 | 1205 |
| **Number of genomes per run** | 24 | 12 |
| **Minimum cost ($USD) per genome** | 229 | 100 |

[a] PacBio library prep kit: SMRTbell prep kit 3.0 (Catalogue no. 102-182-700) (up to 24 samples)
[b] Nanopore library prep kit – Rapid barcoding kit (Catalogue no. SQK-RBK004) (up to 12 plex)

**Table S2.** Cost for Illumina NGS (Singapore based sequencers, Genome Institute of Singapore, A*STAR).

| | **plexWell™96** | **Nextera® XT DNA Sample Preparation Kit (96 Samples)** |
|---|---|---|
| **Library preparation** | 1313 | 2805 |
| **Sequencer- Novogene (2 x 151 bp, 110Gb)** | 1059 | 1059 |
| **Cost per genome (96-plex) ($ USD)** | 25 | 40 |

**Table S3.** Genome assembly and annotation of 8 *Streptomyces* strains

| | **A41733** | **A44034** | **ATCC11455a** | **ATCC21481** | **ATCC23862** | **ATCC31561** | **ATCC31713** | **ATCC31975** |
|---|---|---|---|---|---|---|---|---|
| **Sequence stats** | | | | | | | | |
| Length (bp) | 10516903 | 8284134 | 7110395 | 9265520 | 10548624 | 12115964 | 8794282 | 9566021 |
| Contigs | 8 | 3 | 24 | 7 | 3 | 27 | 186 | 6 |
| GC% | 71.3 | 71.9 | 71.8 | 72.2 | 71 | 70.1 | 72.1 | 70.9 |
| N50 | 10059690 | 8236814 | 5753126 | 7722340 | 10433280 | 1182037 | 70928 | 8753472 |
| Coding density | 85.7 | 83 | 82.1 | 81.6 | 84.1 | 82.4 | 80.3 | 82.4 |
| **Annotation** | | | | | | | | |
| CDSs | 9282 | 8100 | 7099 | 10932 | 11277 | 13086 | 9720 | 9307 |
| Hypotheticals | 2147 | 2679 | 2834 | 6249 | 4742 | 5615 | 5007 | 3620 |
| tRNAs | 77 | 71 | 71 | 75 | 87 | 90 | 72 | 70 |
| tmRNAs | 1 | 1 | 1 | 1 | 2 | 1 | 1 | 1 |
| rRNAs | 21 | 21 | 24 | 24 | 18 | 18 | 3 | 21 |
| ncRNAs | 71 | 26 | 23 | 21 | 32 | 26 | 38 | 53 |
| ncRNA regions | 33 | 24 | 21 | 24 | 27 | 30 | 25 | 23 |
| CRISPR array | 3 | 0 | 0 | 2 | 0 | 1 | 0 | 2 |
| sORFs | 1 | 2 | 1 | 2 | 1 | 3 | 2 | 0 |
| Gaps | 1 | 0 | 0 | 0 | 0 | 0 | 0 | 0 |
| oriCs | 0 | 1 | 1 | 1 | 1 | 1 | 1 | 1 |

**Table S4.** DEREPLICATOR+ metabolite identification

| Strain | Name | Precursor Mass | Retention Time (seconds) | Adduct | Charge |
|---|---|---|---|---|---|
| A44034 | Demethylenenocardamine | 587.337 | 159.184 | M+H | +1 |
| A44034 | Nocardamine_N6-Deoxy / Terragine E | 585.356 | 161.206 | M+H | +1 |
| A44034 | Desferrioxamine E / Nocardamine | 301.18 | 166.356 | M+2H | +2 |
| A44034 | Desferrioxamine E / Nocardamine | 601.35 | 166.11 | M+H | +1 |
| A44034 | Desferrioxamine_X1 | 573.321 | 152.256 | M+H | +1 |
| A44034 | Derhodinyl-streptolydigin;_L-beta1-929 | 487.239 | 230.844 | M+H | +1 |
| A100005 | Banegasine | 205.098 | 121.477 | M+H | +1 |
| A44034 | Demethylenenocardamine | 294.172 | 159.43 | M+2H | +2 |

**Figure S1.** MS/MS spectral comparison of desferrioxamine E and demethylenenorcardamine. Cosine similarity = 0.9748.

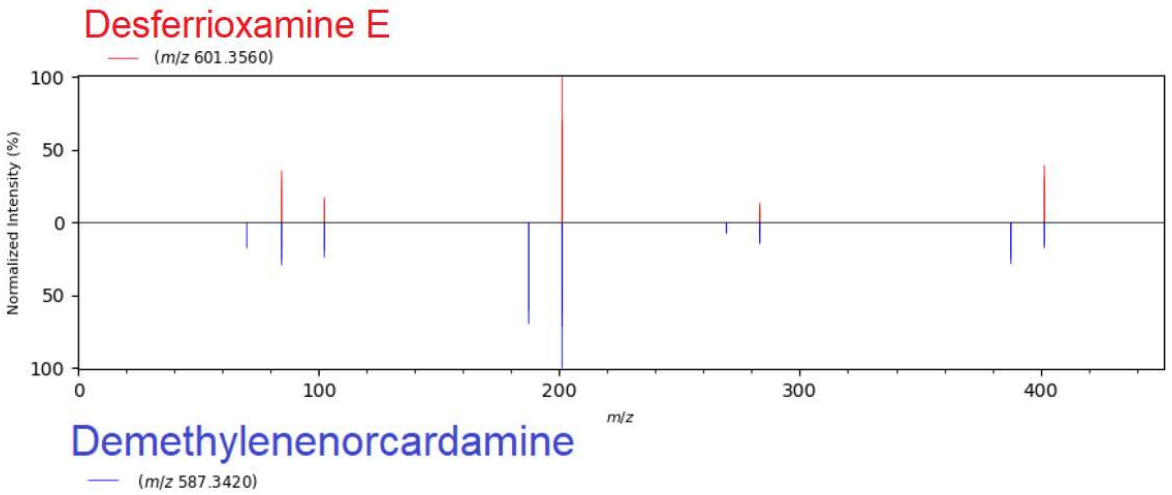


Matching Fragments:

| Desferrioxamine E | Demethylenenorcardamine |
|---|---|
| 84.0788 | 84.0784 |
| 84.0819 | 84.0816 |
| 84.0851 | 84.0848 |
| 102.089 | 102.0886 |
| 102.0925 | 102.0921 |
| 201.1219 | 201.1215 |
| 201.1268 | 201.1264 |
| 201.1317 | 201.1166 |
| 201.1416 | 201.1313 |
| 283.1248 | 283.1244 |
| 283.1307 | 283.1302 |
| 283.1365 | 283.1361 |
| 283.1481 | 283.1419 |
| 401.2523 | 401.238 |

**Figure S2.** MS/MS spectral comparison of desferrioxamine E and Terragine E. Cosine similarity = 0.8360.

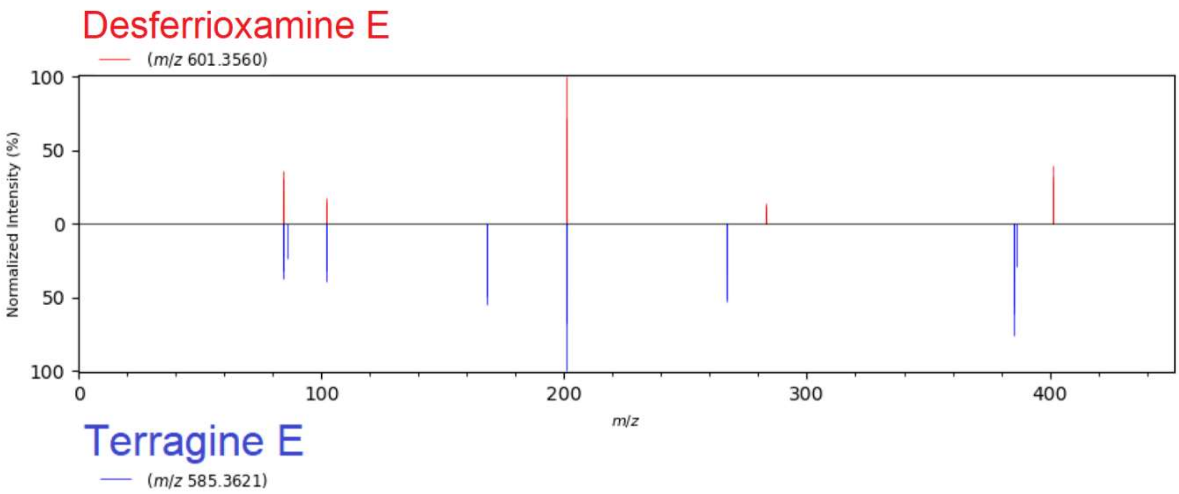


Matching Fragments:

| Desferrioxamine E | Terragine E |
|---|---|
| 84.0788 | 84.0785 |
| 84.0819 | 84.0816 |
| 84.0851 | 84.0848 |
| 102.089 | 102.0887 |
| 102.0925 | 102.0922 |
| 201.117 | 201.1264 |
| 201.1219 | 201.1215 |
| 201.1268 | 201.1166 |
| 201.1317 | 201.1313 |

**Table S5.** Desferrioxamine E gene cluster from A44034, compared to miBIG BGC0001478: desferrioxamine E biosynthetic gene cluster from *Streptomyces sp.* ID38640

| **Query gene** | **Similar protein in NCBI Genbank** | **Percent identity** | **Subject gene** | **Percent Identity** |
|---|---|---|---|---|
| ctg1_5953 | IucA/IucC family siderophore biosynthesis protein | 100.00% | AVV61969.1 | 88.38% |
| ctg1_5954 | GNAT family N-acetyltransferase | 100.00% | AVV61970.1 | 87.83% |
| ctg1_5955 | lysine N(6)-hydroxylase/L-ornithine N(5)-oxygenase family protein | 99.77% | AVV61971.1 | 91.53% |
| ctg1_5956 | aspartate aminotransferase family protein | 100.00% | AVV61972.1 | 89.98% |

**Table S6.** Numbers underlying different pan-genomic 'layers' in Figure 3

| **Layers** | **Total length** | **GC content** | **Percent completion** | **Percent redundancy** | **Number of genes** | **Average gene length** | **Number of genes per kb** | **Singleton gene clusters** | **Number of gene clusters** |
|---|---|---|---|---|---|---|---|---|---|
| **A41733** | 10516903 | 0.713373732 | 100 | 7.042253521 | 9276 | 964.8696636 | 0.882008705 | 1697 | 6335 |
| **A44034** | 8284134 | 0.719431747 | 100 | 5.633802817 | 8102 | 842.9763021 | 0.978014117 | 1318 | 5656 |
| **ATCC11455** | 7110395 | 0.717603874 | 98.5915493 | 4.225352113 | 7089 | 815.4193821 | 0.996991025 | 888 | 4982 |
| **ATCC21481** | 9265520 | 0.722310782 | 95.77464789 | 5.633802817 | 10930 | 686.3280878 | 1.179642373 | 3516 | 5415 |
| **ATCC23862** | 10548624 | 0.70963976 | 98.5915493 | 5.633802817 | 11273 | 782.153464 | 1.068670189 | 3153 | 6269 |
| **ATCC31561** | 12115964 | 0.701494986 | 98.5915493 | 2.816901408 | 13070 | 759.1228003 | 1.078742063 | 4075 | 6517 |
| **ATCC31713** | 8794282 | 0.720826328 | 97.18309859 | 9.85915493 | 9547 | 725.5217346 | 1.085591752 | 2312 | 5425 |
| **ATCC31975** | 9566021 | 0.708668526 | 98.5915493 | 2.816901408 | 9302 | 841.1655558 | 0.972400123 | 2453 | 5588 |
| ***Streptomyces albidoflavus*** | 6841649 | 0.733212709 | 97.18309859 | -1 | 5859 | 1018.515105 | 0.856372491 | 1143 | 4272 |
| ***Streptomyces coelicolor*** | 9054847 | 0.719983673 | 98.5915493 | 1.408450704 | 8159 | 978.7710504 | 0.901064369 | 1667 | 5572 |
| ***Streptomyces griseus*** | 8545929 | 0.722274079 | 94.36619718 | -1 | 7109 | 1055.097763 | 0.831858069 | 1408 | 4998 |
| ***Streptomyces venezuelae*** | 8222198 | 0.724657689 | 98.5915493 | 1.408450704 | 7245 | 1007.157764 | 0.881151244 | 1148 | 5554 |

**Table S7.** BGC classes in the *Streptomyces* strains identified using antiSMASH v6.0 and BiG-SCAPE

| | **PKSI** | **Terpene** | **Others** | **PKS-NRP_Hybrids** | **NRPS** | **RiPPs** | **PKSother** | **Saccharides** |
|---|---|---|---|---|---|---|---|---|
| **A41733** | 4 | 6 | 16 | 2 | 7 | 7 | 3 | 0 |
| **A44034** | 1 | 5 | 7 | 1 | 4 | 5 | 1 | 0 |
| **ATCC11455a** | 2 | 3 | 7 | 0 | 1 | 7 | 2 | 0 |
| **ATCC21481** | 0 | 3 | 18 | 0 | 3 | 5 | 2 | 0 |
| **ATCC23862** | 3 | 6 | 14 | 1 | 5 | 2 | 4 | 0 |
| **ATCC31561** | 0 | 5 | 10 | 1 | 8 | 6 | 3 | 0 |
| **ATCC31713** | 5 | 3 | 10 | 2 | 7 | 2 | 3 | 1 |
| **ATCC31975** | 4 | 4 | 12 | 0 | 3 | 8 | 3 | 0 |
| ***Streptomyces albidoflavus*** | 0 | 3 | 5 | 2 | 5 | 5 | 2 | 0 |
| ***Streptomyces coelicolor*** | 0 | 4 | 8 | 1 | 3 | 5 | 6 | 0 |
| ***Streptomyces griseus*** | 1 | 6 | 12 | 5 | 3 | 8 | 2 | 1 |
| ***Streptomyces venezuelae*** | 0 | 3 | 14 | 1 | 4 | 5 | 3 | 0 |

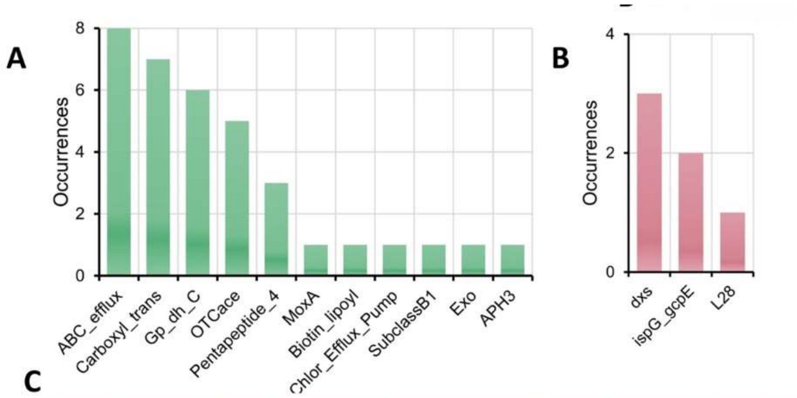


C

| Resistance genes | Function |
|---|---|
| ABC_efflux | ATP-binding cassette (ABC) antibiotic efflux pump [ARO:0010001] |
| APH3 | Streptomycin phosphotransferase |
| Biotin_lipoyl | Biotin-requiring enzyme |
| Carboxyl_trans | Carboxyl transferase domain |
| Chlor_Efflux_Pump | Chloramphenicol efflux pump [ARO:3001212] |
| Exo | Exo beta-lactamase (class a) [ARO:3001213] |
| Gp_dh_C | Glyceraldehyde 3-phosphate dehydrogenase |
| MoxA | MoxA beta-lactamase (class a) [ARO:3001215] |
| OTCace | Aspartate/ornithine carbamoyltransferase |
| Pentapeptide_4 | Pentapeptide repeats (9 copies) |
| SubclassB1 | Subclass B1 (metallo-) beta-lactamase hydrolize penicillins |
| **Core genes** | **Function** |
| dxs | 1-deoxy-D-xylulose-5-phosphate synthase |
| ispG_gcpE | 4-hydroxy-3-methylbut-2-en-1-yl diphosphate synthase |
| L28 | Ribosomal protein bL28 |

**Figure S3.** ARTS annotations for Streptomyces BGCs depicted in Figure 4B and 4C. **(A)** Known resistance genes/mechanisms in BGCs. **(B)** Duplicated core genes in BGCs. C. Abbreviations and names of A and B.
